# CRADLE: A Clinically Robust, Anatomy-Aware Post-Processing Framework for Infant GMA Landmark Tracking in 2D Videos

**DOI:** 10.64898/2026.05.16.725614

**Authors:** Manpreet Kaur, Hamid Abbasi, Angus J.C. McMorland

**Affiliations:** School of Exercise, Sport and Rehabilitation Sciences, Faculty of Science, University of Auckland, Auckland, 1023, New Zealand; Auckland Bioengineering Institute (ABI), University of Auckland, Auckland, 1010, New Zealand; Centre for Brain Research, University of Auckland, Auckland, 1023, New Zealand; Department of Physiology, Faculty of Medical and Health Sciences, University of Auckland, Auckland, 1023, New Zealand

**Keywords:** General movements assessment, infant pose estimation, landmark correction, machine learning, reliable full body tracking

## Abstract

Accurate pose estimation is central to automated infant General Movements Assessment during the fidgety period, when subtle limb movements, particularly at distal joints inform neurodevelopmental risks. Robust 2D pose tracking from handheld videos remains challenging in real-world settings, where occlusion, rapid motions, and visually ambiguous smaller joints frequently compromise anatomical accuracy.

We present CRADLE, a clinically motivated, anatomy-aware post-processing pipeline designed to refine infant 2D movement trajectories across 24-anatomocal landmarks detected by our DeepLabCut-trained model. CRADLE integrates segment-length constraints, velocity-based anomaly detection, anatomically constrained interpolation, and Kalman filtering to correct both large localization failures and subtle persistent joint misplacements without relying primarily on confidence scores.

Evaluations against conventional Confidence-Thresholding using Mean Absolute Error (MAE), ΔMAE, average Percentage of Correct Keypoints, and net keypoint correction rate showed consistently reduced or preserved error while maintaining accurate trajectories, with the strongest gains achieved at clinically important distal joints. Mean improvements reached up to 5 pixels for some smaller distal landmarks, large-magnitude corrections occurred more often than with Confidence-Thresholding, and well-localised joints remained largely unaffected. Positive net correction rates across metacarpophalangeal and metatarsophalangeal distal-landmarks further confirmed a favourable correction-degradation balance.

By improving pose trajectory quality, CRADLE enhances the reliability of downstream movement analysis.

## 1. Introduction

Pose estimation (PE) in video recordings has become a fundamental technique in computer vision, supporting applications such as human activity recognition, sports performance analysis, and person re-identification.^1–3^ While early PE models, such as OpenPose^4^ and Alphapose,^5^ were primarily developed to track adult movements, recent advances in deep learning and transfer learning have enabled their adaptation for infant movement analysis that is useful in clinical applications, including the automated infant General Movements Assessment (GMA). GMA is a validated clinical tool for assessing spontaneous infant movements during early postnatal development. Fidgety movements (FMs) are an important subtype of GMA, typically emerging between 9 and 16 weeks post-term, that are known to be strongly associated with later neurodevelopmental disorders such as cerebral palsy (CP) when absent or abnormal.

Traditionally, the GMA is conducted manually by trained clinicians who visually evaluate infant videos.^6^ While the manual GMA predicts CP with around 95% sensitivity and specificity,^7^ it is time- and resource-intensive, requires expert training, and has limited scalability. To address these limitations, automated GMA approaches have been developed to improve the speed and scalability of the assessment.^8–12^ These systems typically derive features from PE-based anatomical keypoints and trajectories to quantify clinically relevant information such as movement complexity, variability and fluidity, which are then used to classify movement quality and neurodevelopmental risk.^8,13^

Despite progress, accurate infant PE from 2D recordings remains highly challenging due to issues such as frequent landmark occlusion, unstructured and rapid limb movements, and limited pose diversity in PE training data.^14,15^ These factors often lead to inaccurate keypoint tracking, especially for smaller and more visually changeable distal joints such as metacarpophalangeal (MCP) and metatarsophalangeal (MTP) joints in the hands and feet.^16^ Fine, continuous, rotational movements of the wrist that require tracking of these distal joints are thought to carry important information for an accurate GMA determination.^17^ Errors in the tracking of such landmarks can distort movement trajectories and degrade the feature quality, ultimately affecting the performance and generalizability of the automated GMA systems and reducing their clinical reliability.

A commonly used post-processing strategy to correct infant PEs is to apply a dynamic confidence threshold (e.g., 90% of the average confidence for a given keypoint) to remove low-confidence predictions, followed by interpolation to fill the resulting gaps.^18,19^ This approach assumes that lower confidence scores correspond to inaccurate predictions.

However, confidence values reflect model certainty rather than anatomical correctness. Consequently, confidence-based post-processing may discard visually correct keypoints when confidence is low (e.g., due to occlusion or rapid movement) and may retain anatomically incorrect keypoints when confidence is high. While the extent and impact of these limitations in infant pose predictions remain unexplored, a priori there is a need for more robust post-processing techniques that move beyond confidence scores to generate anatomically consistent and clinically reliable pose data, particularly for downstream tasks such as automated GMA, where precision is critical.

In this paper, we propose CRADLE (clinically-robust anatomy-aware distal landmark enhancement), a novel, multi-stage post-processing pipeline to detect and correct keypoint errors in 2D infant pose trajectories estimated by a PE model (i.e., a DeepLabCut(DLC)^20^ trained ResNet152). CRADLE is capable of addressing distinct types of keypoint errors by integrating segment length- and velocity-based analysis, unsupervised clustering, Kalman filtering,^17,21^ and interpolation with anatomical constraints. To our best knowledge, this is the first dedicated post-processing approach aimed at correcting PE errors in infant movement data, a critical yet overlooked aspect in the existing literature. While most prior efforts have focused on improving core model architectures or downstream classification methods, our work improves the quality of the input pose data itself. By correcting keypoint tracking inaccuracies, especially in clinically sensitive joints such as MCPs and MTPs, our CRADLE enhances the reliability of anatomical consistency and temporal continuity of pose trajectories, an approach that is highly useful in infant automated GMA and other clinical movement analyses.

## 2. Participants and Data Collection

### 2.1 Ethics

All study procedures were approved by the Auckland Health Research Ethics Committee (AHREC-000146). Parents or caregivers were fully informed about the purpose of the study, video-recording procedures and subsequent data use. Written informed consent was obtained for each infant participant.

### 2.2 Recording Procedures

Clinically recorded videos were obtained from the Child Development Centre of the Waikato District Health Board (DHB), New Zealand. For this study, we selected a diverse subset of 12 full-term infants (8 with clinically confirmed normal FMs), with a mean gestational age of 31.25 weeks (SD: 5.46), mean birth weight of 1619 grams (SD: 989.80), and mean corrected age of 91.08 days (SD: 8.95) at the time of recording. Infants were selected from a large clinical database to ensure diversity in movement patterns, partial occlusions, skin tones, clothing, background settings, camera angles and lighting conditions to promote generalizability. All recordings were captured during the infants’ FM period (i.e., age range: 9–16 weeks post-term) while the infants were awake, calm, and in a natural behavioural state. Each infant was placed in a supine position on a plain-coloured linen sheet over a comfortable mat and wore only a nappy. Video recordings were conducted using standard iPads (MQDT2X/A: 12-megapixel camera with 4K HD video and a MD367X/A: 3^rd^ generation iPad with 5-megapixel 1080p HD camera) at a resolution of 1920 x 1080 pixels and a frame rate of 29.97 frames/second. Each video had a duration ranging from 1.5 to 3 minutes. Video data were trimmed using Adobe Premiere Pro 2021 to remove the initial and end video portions where the infant was out of frame, ensuring recordings contained the infant within the center of the frame. This step helped to also crop excessive background while preserving the original video quality.

## 3. Methods

### 3.1. Pose Estimation

We trained ResNet-152 using the DLC environment,^20^ for markerless PE to extract 2D trajectories of 24 anatomical keypoints (right & left eyes, nose, sternum, 5 landmarks per upper limb: shoulder, elbow, wrist, little finger MCP and index finger MCP, and 5 landmarks per lower limb: ASIS, knee, ankle, little toe MTP and big toe MTP)^22^ in 12 infant videos. The repeated hold-out procedure was performed five times to train 15 independent models. Each model was trained for 700,000 iterations on our custom-labelled dataset comprising 100 annotated frames per video (total 8700 frames).^23^

In order to rigorously evaluate the robustness of the CRADLE pipeline in this paper, we selected the model with the lowest cross-validation accuracy (i.e., 98.70%) for generating the initial pose trajectories used in this study. This selection ensures that the input data includes a realistic proportion of keypoint inaccuracies, particularly in joints such as hands and feet. For quantitative evaluation, ground truth trajectories were generated for all the frames in 12 videos by first applying the best-performing model (accuracy: 99.30%), followed by manual correction of its outputs, including challenging cases such as occlusion and motion blur, where keypoints were annotated based on the best possible estimate of their true locations. This strategy was more efficient than annotating frames from scratch and enabled frame-wise evaluation using performance metrics.

### 3.2. Proposed Pipeline: CRADLE

The flowchart of our proposed, CRADLE, post-processing approach for automatically detecting and correcting erroneous keypoints in the 2D pose trajectories (generated through any deep-learning-based method—DLC in our case), is illustrated in **Figure 1**. The detailed pseudocode of the approach is provided in the supporting material (**Table S1**).

**Figure 1.**
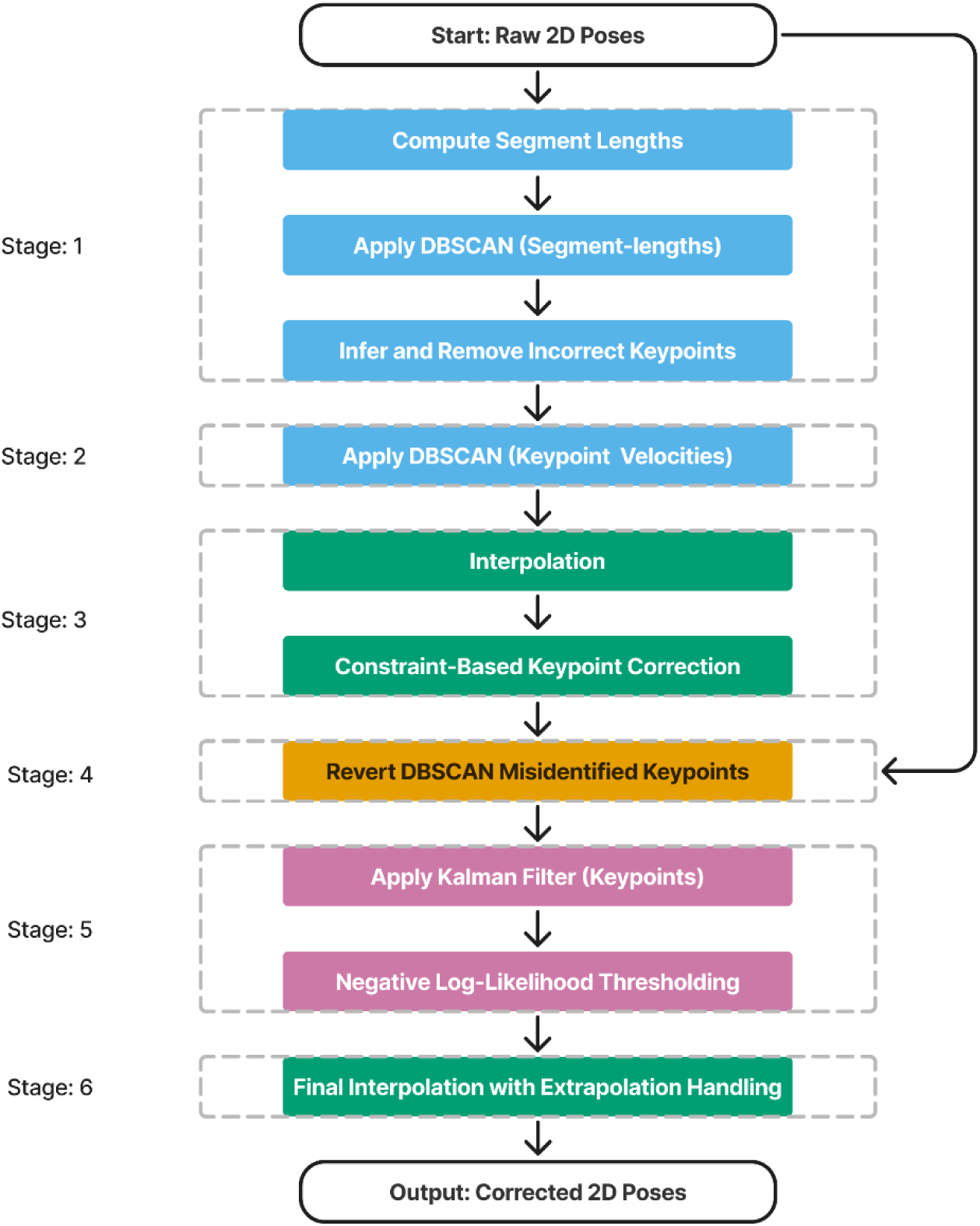
Flowchart of the CRADLE post-processing pipeline with six sequential stages. The method comprises six sequential stages: (1) segment length-based error detection, (2) velocity-based error detection, (3) anatomically constrained interpolation, (4) reversion of misclassified keypoints, (5) Kalman filter-based refinement, and (6) final interpolation with extrapolation handling.

The first four stages primarily target large, unrealistic errors, including anatomically inconsistent limb configurations, extreme segment distortions, and abrupt spatial discontinuities (e.g., mislocalization of MCPs over MTPs and vice versa). In contrast, the final two stages refine subtle yet clinically important inaccuracies that persist after initial corrections, such as keypoints’ minor displacements caused by soft occlusions. Each stage is described in detail below.

#### 3.2.1. Segment Length-based Error Detection

To identify erroneous keypoints, we computed 29 segment lengths per frame using 24 anatomical keypoints (see **Figure 2**). Each segment length, numbered 1–29, corresponds to the Euclidean distance between anatomically adjacent keypoints. The underlying assumption was that keypoints mislocalized for a block of frames would cause abrupt or inconsistent changes in at least one of their connected segments over time. From here on, for simplicity, we refer to these anatomical segment lengths as ‘segment lengths’.

**Figure 2.**
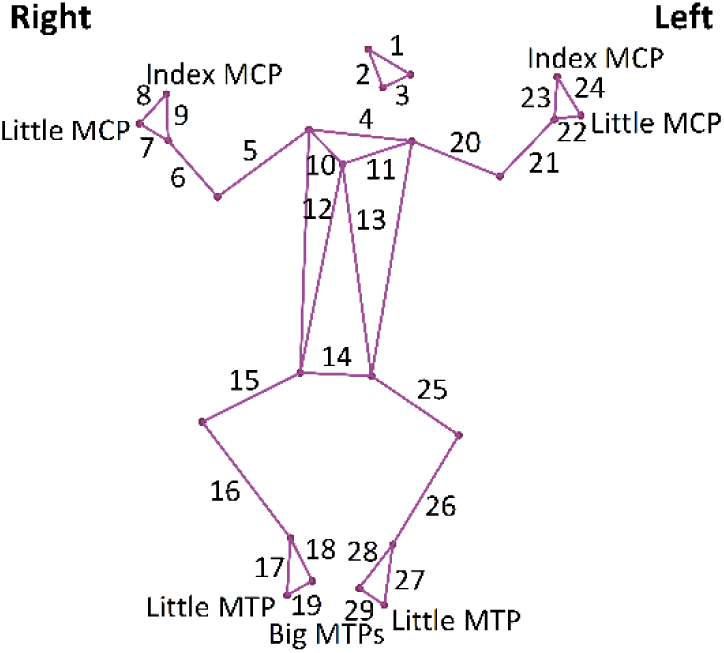
24 anatomical keypoints and 29 segments connecting them.

We applied Density-Based Spatial Clustering of Applications with Noise (DBSCAN),^24^ an unsupervised density-based clustering algorithm, independently to the time series of each segment length. DBSCAN identifies clusters by grouping data points (here, segment lengths across frames), that are spatially close in the feature space, using two parameters: the neighbourhood radius (*ε*) and the minimum number of points (*minpts*) required to form a cluster. The algorithm begins at a random point and expands clusters by including neighbouring points that satisfy the density criterion, that is, having at least *minpts* points within the *ε*-neighbourhood. The values of *ε*, 0.5 for face, 0.25 for trunk, and 0.6 for limb keypoints, and *minpts,* 20 for body, and 40 for face keypoints, were determined empirically through systematic sensitivity analysis by using multiple combinations of *ε* (0.1–0.8) and *minpts* (10–60) across representative videos. For each parameter combination, the resulting cluster structure was visually inspected to assess the separation between dense clusters and sparse outliers. Then, parameters were selected based on their ability to form a stable high-density cluster corresponding to possible consistent segment lengths, while reliably isolating abrupt fluctuations and erroneous values. Given region-specific movement variability, different parameter settings were adopted for face, trunk, and limb segments. Points that do not belong to any cluster were considered outliers (or errors) and were flagged for further inspection as potentially incorrect segment lengths.

To infer incorrect keypoints from the flagged segment lengths, we implemented a set of hierarchical logical rules (detailed in Supporting material Table S1). These rules systematically evaluate the segment length abnormalities in a proximal-to-distal manner, based on the assumption that proximal joints (e.g., shoulders, ASIS) are more stable and can be used to guide the inference of more mobile distal joints (e.g., wrists, ankles, MCPs, MTPs). Inference logic was defined based on the number and configuration of affected adjacent keypoints, and errors were categorised into three main types:

##### Single-keypoint errors

These involve one keypoint connected to two flagged segments while adjacent keypoints are connected to segments marked as correct. Such cases were handled using simple logical rules. For example (Figure 2), if both segments, shoulder–elbow and elbow–wrist, are flagged, but the shoulder and wrist keypoints are correct, then elbow keypoint is inferred as incorrect. This logic was encoded as:

If [(seg_4 & seg_10 & seg_9 & seg_7 == C) & (seg_6 or seg_5 == I)]: → Rt_elbow

##### Two-keypoint errors

These typically occur in triangular configurations, such as the hand or foot, where two adjacent segments (e.g., wrist–index MCP and wrist–little MCP) are flagged. To resolve complex ambiguities regarding whether the error occurred at the wrist, or both MCPs (see Figure 2), a specific rule was applied. For example,

If [(seg_6 == C) and (seg_9 & seg_7 == I)]: → Both Rt_MCPs

##### Three-keypoint errors

These involve all three keypoints forming an anatomical triangle (e.g., wrist and both MCPs) connected by all three flagged segments. In such cases, all three keypoints were classified as incorrect based on rules, for example:

If (seg_5 & seg_6 & seg_7 == I): → Rt_wrist and both Rt_MCPs

These logical rules were applied symmetrically across left and right limbs to preserve anatomical consistency throughout the correction process. Subsequently, all keypoints identified as incorrect were replaced by NaN values to indicate missing or invalid keypoints.

#### 3.2.2. Velocity-based Error Detection

To detect remaining mislocalized keypoints not identified by segment length-based filtering, we focused on 12 distal keypoints (bilateral wrists, ankles, MCPs and MTPs) from stage 1. Frame-to-frame 2D keypoint velocities were computed, and DBSCAN (*ε* = 30, *minpts* = 20) was applied to identify anomalous motion patterns and flag corresponding keypoints. A larger neighbourhood radius was used at this stage to account for the initial removal of large localization errors and to avoid marking normal small movement variations as outliers, which can occur when smaller neighbourhood values produce excessive clustering. Parameter tuning was carried out by testing *ε* values between 10 and 50, and *ε* = 30 was selected based on visual inspection to retain normal motion while consistently identifying abrupt velocity jumps. To further refine detection, a ±2-frame window around each flagged keypoint was then examined. If a keypoint’s velocity was near zero despite neighbouring keypoints being flagged, we extended the flag to these adjacent frames, indicating temporally consistent but erroneous localizations, where the keypoint remained in an incorrect position across several continuous frames. All keypoints flagged in this stage were replaced by NaN values prior to the next correction stage.

#### 3.2.3. Anatomically Constrained Interpolation

Following the previous filtering, modified Akima interpolation^25^ was applied to fill in missing NaN values. Further, to prevent unrealistic segment length deviation from expected anatomical proportions, during interpolation over large gaps (e.g., ≥ 2–3 seconds), especially for distal joints (e.g., MCPs and MTPs) that move and rotate more extensively relative to proximal joints, anatomical constraints were imposed on hand and foot segment lengths. Specifically, the 99^th^ percentile of each segment length from the pre-interpolation data was computed, where the majority of keypoints were correct, and this value was used as the maximum allowable segment length (hereafter referred to as a threshold) for further processing.

When an interpolated distal keypoint violated more than one connected segment length constraint (e.g., both wrist–MCP and MCP–MCP thresholds), two intermediate positions were computed. Each position was obtained as the point along the segment vector, shortened to the threshold length. The final corrected location was then defined as the midpoint of these two intermediate positions. In contrast, when both distal keypoints were displaced (e.g., both wrist–MCP distances exceeded their thresholds), each keypoint was corrected independently by constraining it to the threshold distance along the same direction vector from the proximal joint (wrist/ankle). In this case, no averaging was applied, as each correction involved only a single segment-length constraint. Through these processes, keypoints’ positions were adjusted so that the segment lengths remained within anatomically realistic limits, while maintaining anatomically consistent trajectories.

#### 3.2.4. Reversion of Misclassified Keypoints

To prevent overcorrection resulting from DBSCAN’s parameter sensitivity, a reversion step was implemented that selectively replaced interpolated keypoint positions with restored keypoints from the original PE output, when either of two conditions was met:

a. If the interpolated keypoint was within 10% of the maximum torso length (used as spatial threshold) from the PE keypoint, it was reverted to retain the model’s accurate predictions.
b. If the original segment length (from the PE model) was within the 99^th^ percentile of segment lengths computed from pre-interpolation data (i.e. output of Stage 2), and the interpolated segment exceeded this range, the original keypoint was restored.

#### 3.2.5. Kalman Filter-based Refinement

To further enhance the spatial precision of keypoints with potential slight displacements from their true locations due to occlusions or rapid joint movement, each keypoint trajectory was refined using a Kalman filter. The negative log-likelihood of observed positions was computed. Frames with a likelihood below a threshold (-25), selected empirically, were considered uncertain and temporarily removed by replacing them with NaN values, resulting in short gaps in the trajectories.

#### 3.2.6. Final Interpolation with Extrapolation Handling

Gaps introduced by the Kalman filter (in Step 5) were filled using modified Akima interpolation to maintain continuous keypoint trajectories, completing the final correction step. However, interpolation at the temporal boundaries (start/end of the series) can be unreliable due to missing reference points on one side; therefore, any boundary frames flagged as incorrect during Stage 1 were excluded.

Our methodology of implementing these six stages was designed to help integrate spatial, temporal, and anatomical information, obtained through sequential frames, aiming to improve the predicted anatomical location consistency and temporal continuity of pose trajectories for 2D video-based infant PE.

### 3.2. Baseline: Confidence-Thresholding

To benchmark the performance of our CRADLE pipeline, we also implemented an alternative, widely used Confidence-Thresholding method. For each keypoint, we calculated its average likelihood across all the frames and set a dynamic threshold at 90% of its value.^7,23^ Keypoints with likelihood below their respective thresholds were removed and replaced with NaN values. The resulting gaps were then interpolated using modified Akima interpolation, and boundary extrapolation was handled as in Stage 6 of our pipeline, ensuring a fair comparison across identical frames.

### 3.3. Performance Measures

Both quantitative and qualitative comparisons were made between the CRADLE pipeline and the alternative Confidence-Thresholding approach. For quantitative assessment, four metrics were used:

#### 3.3.1. Mean Absolute Error (MAE)

MAE was calculated for each keypoint as the Euclidean distance between ground truth keypoints and the following three datasets, on a frame-by-frame basis: (i) raw PE output, (ii) keypoints after applying Confidence-Thresholding, and (iii) keypoints after applying the CRADLE method.

#### 3.3.2. ΔMAE

To evaluate the overall effectiveness of post-processing methods (CRADLE and Confidence-Thresholding), we calculated ΔMAE for each keypoint, defined as the average MAE (across frames) ‘before’ minus ‘after’ post-processing. A 95% confidence interval of ΔMAE across all infants was then calculated. Positive ΔMAE values indicate a reduction in keypoint localization error, whereas negative values reflect increased error after post-processing. To further assess the performance of post-processing methods at the frame level, the distribution of frame-wise error changes (in pixels) across 12 distal joints was analysed. For each frame, a ΔMAE was computed across infants and keypoints. ΔMAE-frame frequencies were then binned and summarized to characterize the distribution of per-frame error changes for each method.

In addition, to examine whether the error improvement magnitude depended on the MAE before post-processing (computed from the raw PE output), the mean MAE of each distal keypoint (averaged across test videos) was plotted against its corresponding ΔMAE.

#### 3.3.3. Average Percentage of Correct Keypoints (aPCK)

Keypoint localization accuracy was also evaluated using aPCK at thresholds of 5%, 7%, and 10% of maximum torso length, (aPCK @0.05, aPCK @0.07, and aPCK @0.1). A keypoint was considered correct if its distance from the ground truth location was within the corresponding threshold radius. As with MAE, aPCK was computed before and after post-processing (CRADLE and Confidence-Thresholding).

#### 3.3.4. Net Keypoint Correction Rate

To assess frame-wise correction effectiveness of each keypoint, we quantified the net change in keypoint correctness using the same aPCK thresholds. For each keypoint, frames were categorized as recovered, where an initially incorrect keypoint became correct after post-processing, or worsened, where an initially correct keypoint became incorrect. The net correction rate was defined as the difference between recovered and worsened frames, normalized by the total number of frames and scaled per 10,000 to improve interpretability due to the small magnitude of absolute changes, such that,

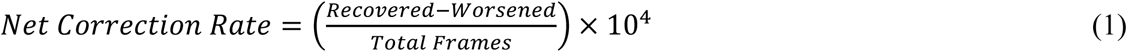

While both post-processing approaches (CRADLE and Confidence-Thresholding) were applied to all 24 keypoints (Figure 2), our evaluation primarily focused on anatomically distal keypoints (hands and feet), which are known to be more challenging for PE models due to higher mobility and frequent occlusions compared to other body landmarks.

### 3.4. Processing Environment

All experiments were conducted on a Windows 10 system with a 12^th^ Generation Intel ® Core ^TM^ i5-1235U processor, 16GB RAM, and a 64-bit Operating System. The implementation was done in Python 3.10.9 using Jupyter Notebook (v6.5.2).

## 4. Results

We assessed the performance of CRADLE, a novel post-processing algorithm for PE data, against an alternative, commonly used algorithm, Confidence-Thresholding.

### 4.1. Quantitative Assessment

**Figure 3–7** show infant- and keypoint-level results for each hand and foot keypoints using MAE, ΔMAE, and aPCK, and **Table 1** reports Net Keypoint Correction Rate. Results are presented for three trajectory types: (i) PE predictions (Before); (ii) Confidence-Thresholding; and (iii) CRADLE’s output. Summary statistics averaged across all 12 infants are provided in supporting material **Table S3–S5**.

**Figure 3.**
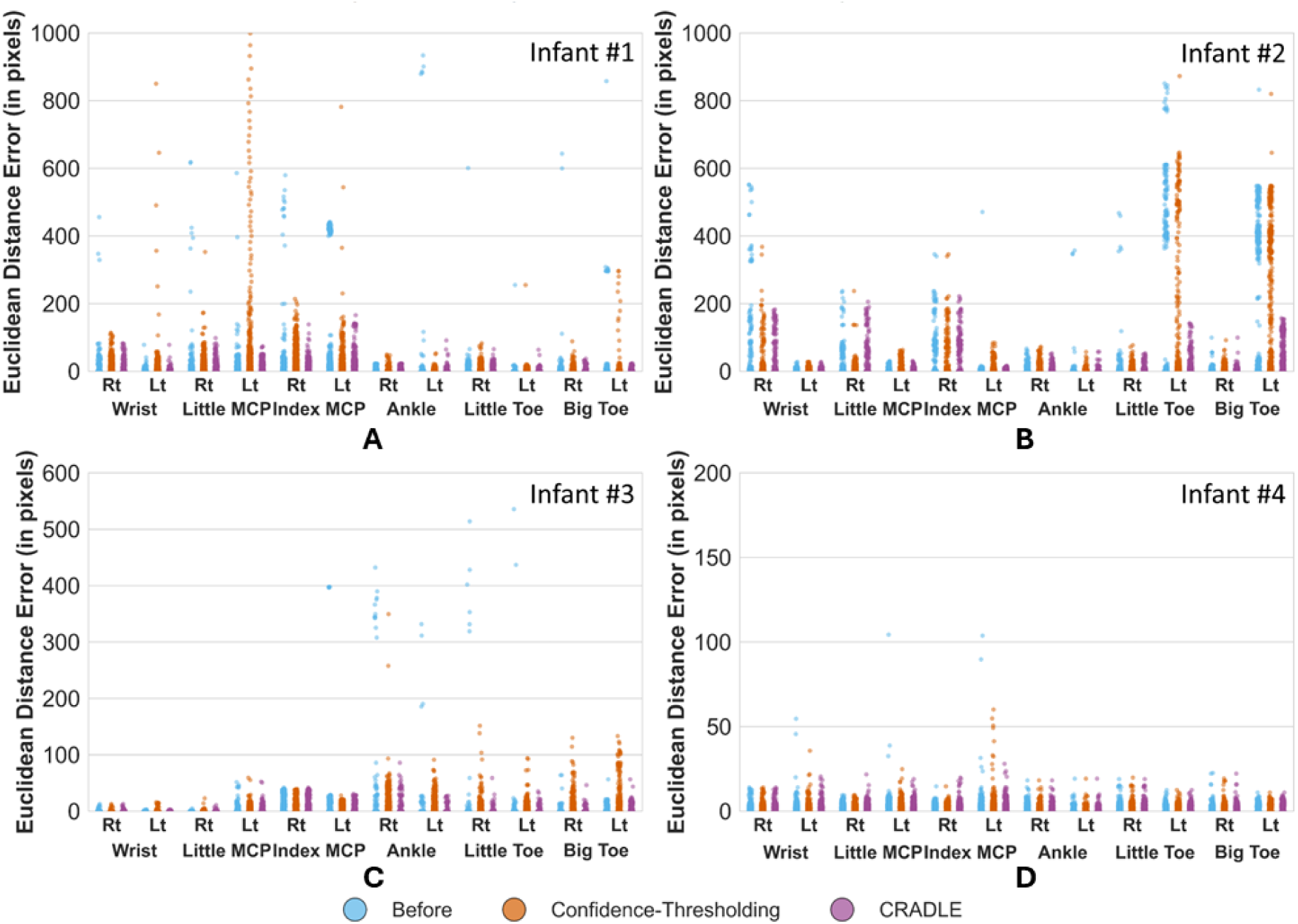
Mean absolute error (MAE) distributions for four representative infants (#1–#4). Blue, orange, and purple markers indicate the Before, Confidence-Thresholding, and CRADLE methods, respectively. Each plot (A–D) shows per-frame MAE (y-axis) for individual keypoints (x-axis) and each dot corresponds to a single frame.

**Figure 4.**
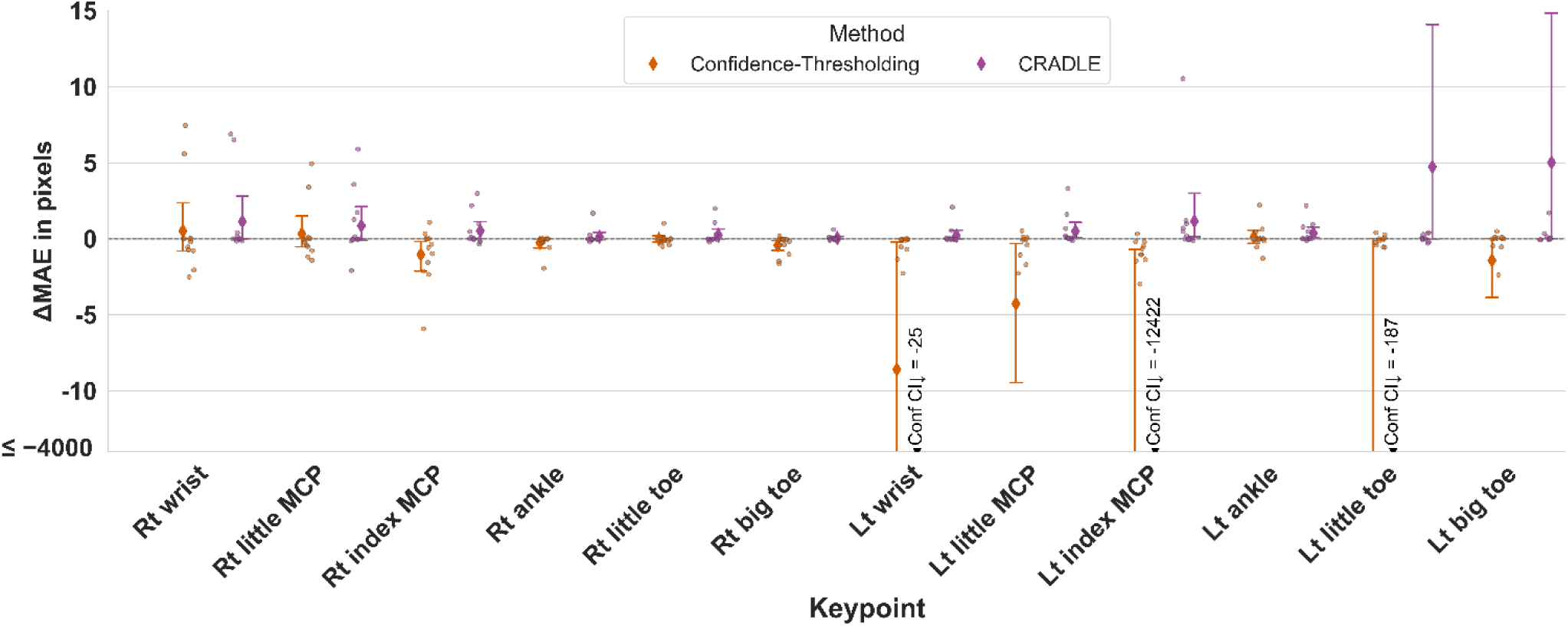
ΔMAE (MAE_before_ − MAE_after_) comparison between Confidence-Thresholding (orange) and CRADLE method (purple) across 12 infants for each keypoint. Blue, orange, and purple markers indicate the Before, Confidence-Thresholding, and CRADLE methods, respectively. Each dot represents one infant, with vertical bars showing 95% confidence intervals. Positive values indicate improved accuracy after correction.

**Figure 5.**
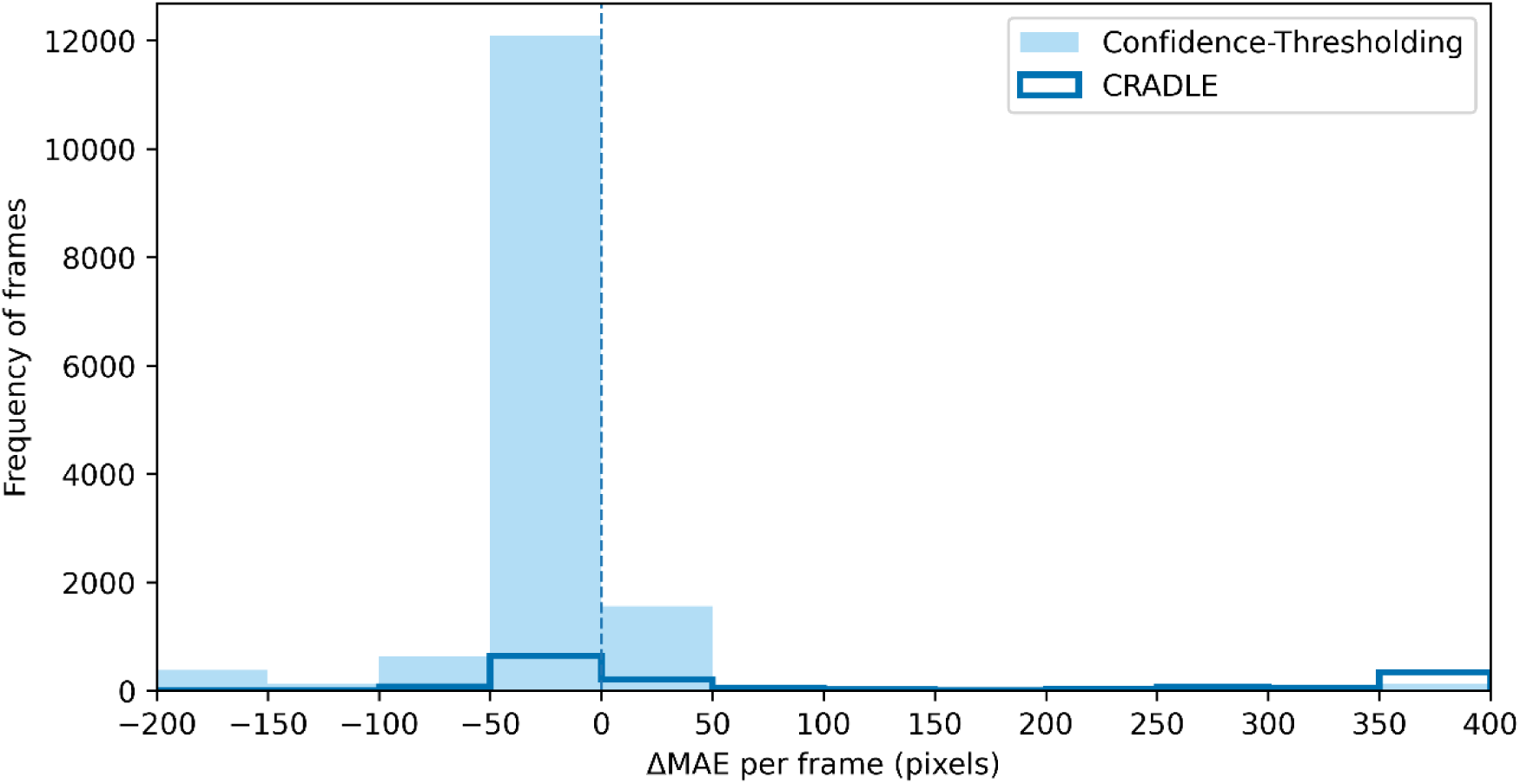
Distribution of frame-wise ΔMAE (MAE_before_ – MAE_after_) across all distal keypoints and test videos for Confidence-Thresholding (light blue) and CRADLE method (dark blue outline). Each bin represents the frequency of frames within a given ΔMAE range. The vertical dashed line indicates ΔMAE = 0, where positive values correspond to reduction in localization error after post-processing and negative values indicate an increase in error.

**Figure 6.**
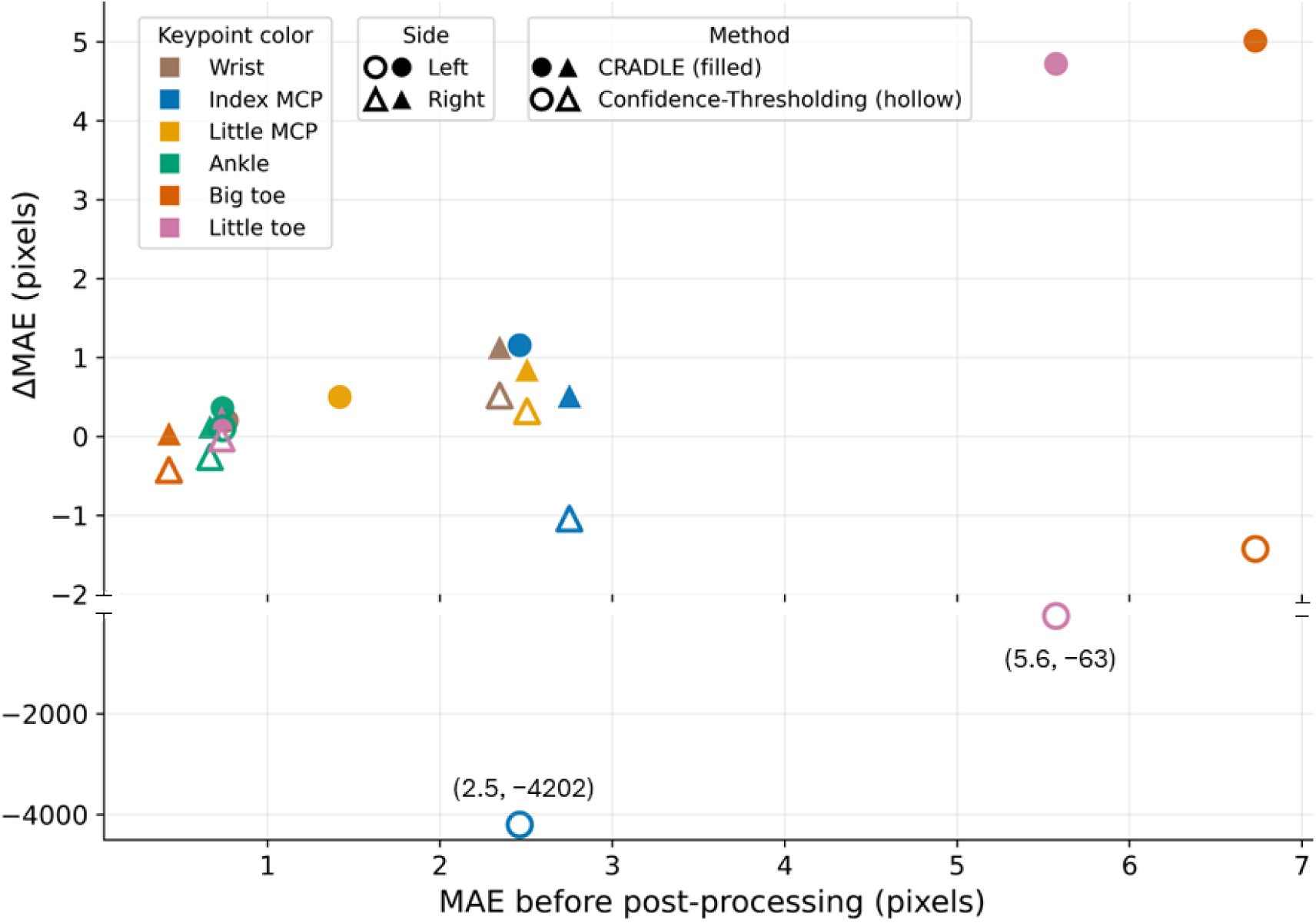
Relationship between baseline keypoint error (MAE_before_) and ΔMAE (MAE_before_ − MAE_after_) for the 12 distal keypoints. Filled markers represent CRADLE, and hollow markers represent Confidence-Thresholding; colours indicate keypoint type and marker shape distinguishes left and right sides. Each point corresponds to the mean value across test videos. Positive ΔMAE values indicate improved localization following post-processing.

**Figure 7.**
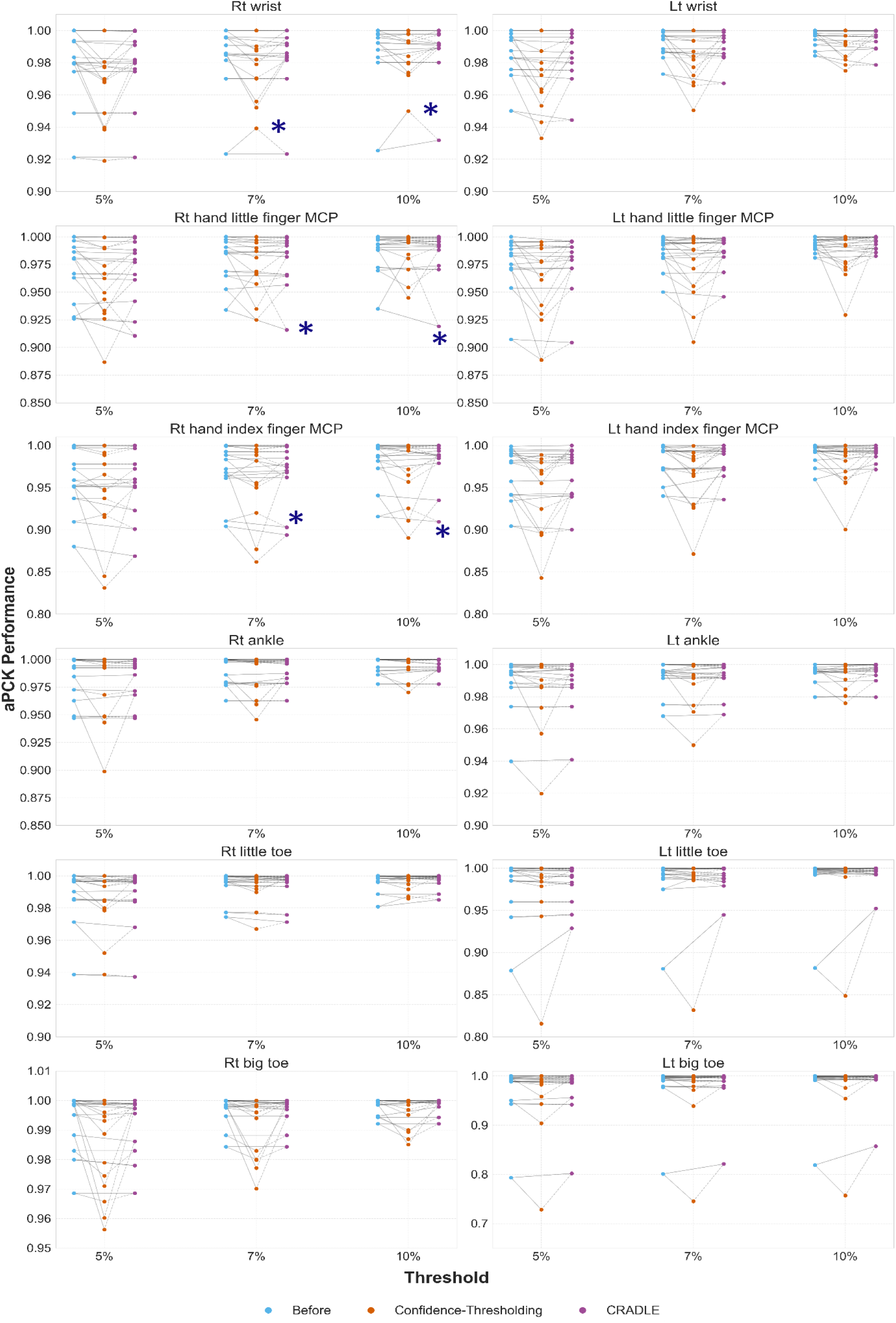
aPCK values for hand and foot keypoints across three thresholds (5%, 7%, 10%) for all 12 infants. Blue, orange, and purple markers indicate the Before, Confidence-Thresholding, and CRADLE methods, respectively. Each point represents one infant. The asterisk (*) highlights Infant #2, corresponding to one of the infants presented in Figure 3, to illustrate how performance changes for the same infant under the aPCK metric.

**Table 1.**
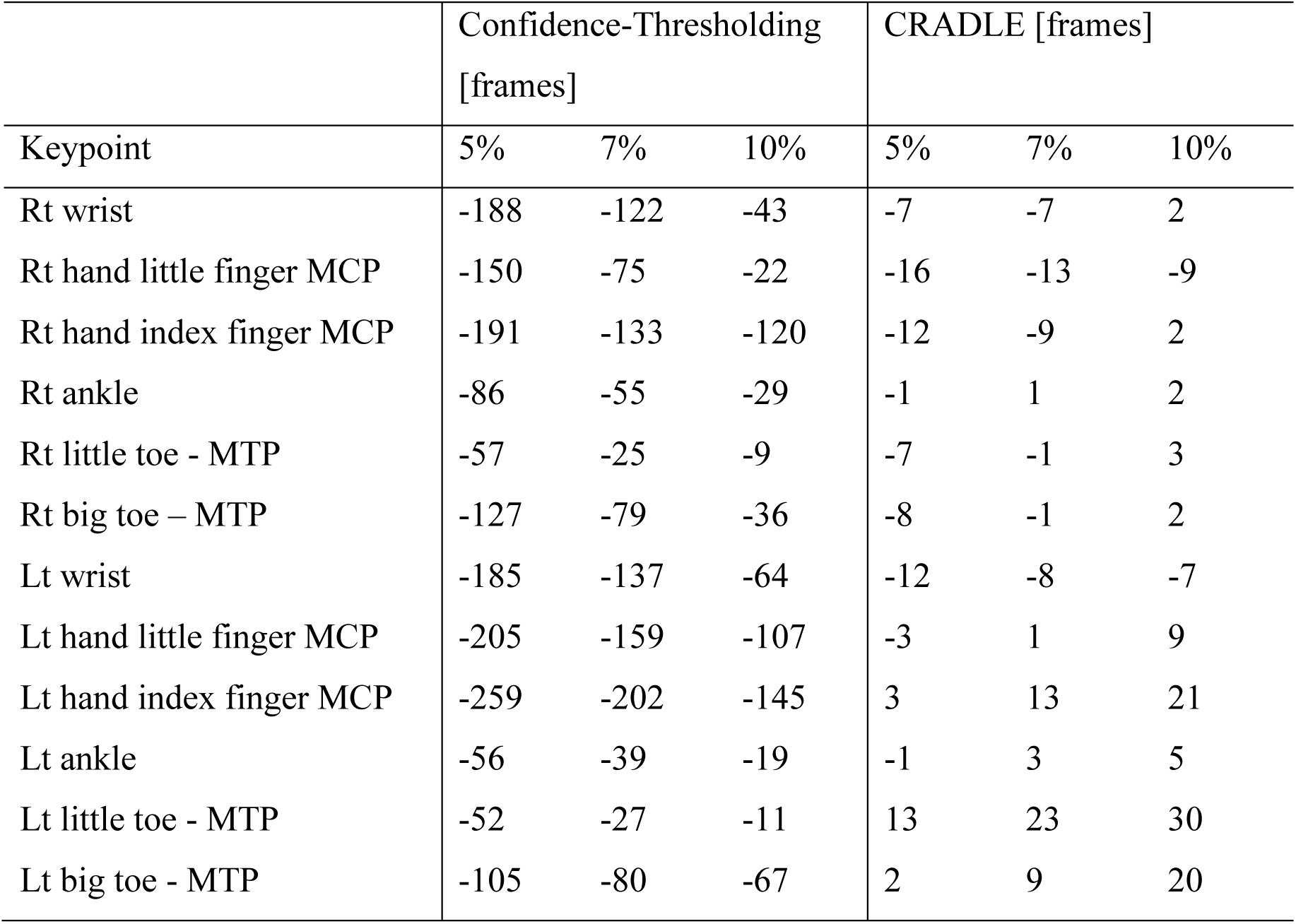
Net Keypoint Correction Rate (per 10,000 frames) for hand and foot keypoints across three thresholds (5%, 7%, 10%) for 12 infants.

#### 4.1.1. MAE

**Figure 3** shows MAE distributions for four representative infants (A–D), selected to illustrate the range of performance patterns observed across the cohort. These infants capture distinct scenarios (detailed in the Discussion Section). Similar patterns were observed for the remaining infants, and are documented in the supporting material **Figure S2**. In Figure 3, the CRADLE method reduced large errors or maintained error levels comparable to the ‘Before’ predictions for all infants. In contrast, Confidence-Thresholding resulted in higher MAE across several keypoints for Infant #1 and #3, including the left little MCP and big toes. Similarly, for Infant #2, Confidence-Thresholding showed increased MAE for the left little and big toes, while the CRADLE method achieved lower MAE across most keypoints, although some (e.g., right little MCPs) showed only small improvement. For Infant #4, where original PE predictions were already accurate, both methods produced low MAE with negligible differences.

#### 4.1.2. ΔMAE

**Figure 4** presents the distributions of ΔMAE in pixels (px) across infants for each keypoint, shown as strip plots where each dot represents one infant and vertical lines indicate 95% confidence intervals (CIs). For several keypoints, such as left wrist, and left index MCP, left little MTP, the Confidence-Thresholding method produced extreme errors, resulting in unusually elongated 95% CIs (see section Qualitative Assessment for an explanation of how this arises). To enhance the visual interpretability for most cases, the y-axis in Figure 4 was restricted to –10 to +15 px. Extremely large negative values (≤ –4000 px) were represented at the lower bound, with actual values annotated on the corresponding vertical lines, which consequently truncated CI lines for these extreme keypoints. Full statistics, including means and standard deviations (SDs), are provided in supporting material **Table S3**.

Figure 4 shows that the CRADLE method consistently improved performance across nearly all keypoints, with positive ΔMAE values and reduced variability. Notably, left little and big toes improved by 4.72 px (SD = 14.83) and 5.01 px (SD = 13.26), respectively, while hand joints, including the left index MCP and right wrist, improved by 1.16 px and 1.13 px. In contrast, Confidence-Thresholding produced unstable and sometimes severely degraded outcomes, increasing MAE for the left index MCP by 4202.17 px (SD = 23,827.05) and the left little toe by 63.32 px (SD = 524.89) (Table S3).

**Figure 5** further illustrates the distribution of frame-wise ΔMAE across all 12 distal keypoints. Confidence-Thresholding produced a large proportion of frames in the negative ΔMAE range (0 to −50 px), indicating frequent increases in localization error after post-processing. In contrast, CRADLE resulted in substantially fewer frames within this negative region. While Confidence-Thresholding produced moderate positive changes (up to ∼50 px) across a higher number of frames, CRADLE exhibited fewer but larger positive ΔMAE shifts, including values in the 350–400 px range, reflecting correction of larger localization errors.

**Figure 6** presents the relationship between MAE before post-processing and ΔMAE. Keypoints with higher MAE before post-processing (e.g., left little and big toes) showed larger positive ΔMAE following CRADLE, whereas keypoints with low MAE before post-processing (e.g., right big toe, and ankle) remained close to zero. Confidence-Thresholding showed inconsistent behaviour, including a significant reduction (extreme negative) in ΔMAE for certain keypoints (left little and big toes) despite moderate MAE before post-processing.

#### 4.1.3. aPCK

**Figure 7** shows aPCK for hand and foot keypoints at three thresholds (5%, 7%, 10%). Each subplot corresponds to one keypoint location, with columns representing performance for the original predictions (Before), Confidence-Thresholding, nd CRADLE method. Each point corresponds to one infant. The average summary statistics are provided in supporting material **Table S4**.

Across all thresholds, the CRADLE method maintained or slightly improved aPCK compared to the Before predictions. Notable improvements were observed for the left little toe, average across 12 infants, increasing from 0.978, 0.984, 0.988 (Before) to 0.981, 0.989, and 0.993 at 5%, 7%, and 10% thresholds, respectively. The left big toe increased from 0.970 to 0.986 at the 10% threshold (Table S3).

By contrast, Confidence-Thresholding often reduced aPCK for an infant highlighted with * in Figure 7, which corresponds to same infant #2 in Figure 3. For this infant, the left index MCP decreased from 0.967 (Before) to 0.939, 0.958, and 0.973 across the different thresholds. Similar reductions were observed for the right index MCP (0.957 → 0.939 at 5%) and the right wrist (0.979 → 0.966 at 5%), with the largest differences at the strictest, 5% threshold.

#### 4.1.5. Net Keypoint Correction Rate

**Table 1** shows the Net Keypoint Correction Rate per 10,000 frames. Positive values indicate a net improvement (i.e., more errors brought below the threshold than raised above it), while negative values indicate net degradation in keypoint localization. Results in Table 1 indicate that the CRADLE method consistently achieved higher or less negative net correction rates across most keypoints and thresholds compared to Confidence-Thresholding, which yielded higher negative values across all keypoints. For example, at the left index finger MCP, our CRADLE method corrected 3, 13, and 21 keypoints per 10,000 frames at the 5%, 7%, and 10% thresholds, respectively, whereas the Confidence-Thresholding method produced net changes of -259, -202, and -145 at the corresponding thresholds. Similar trends were observed for MTPs and wrists (Table 1).

### 4.2. Qualitative Assessment

**Figure 8** and **9**, and **Table 2** present representative visualizations of keypoint trajectories overlaid on video frames to illustrate the performance of each post-processing method. In these plots, blue lines represent original PE model outputs, orange lines correspond to the Confidence-Thresholding method, gray lines denote intermediate incorrect outputs from the CRADLE pipeline, and purple lines indicate the final output from the CRADLE method. Black Xs mark keypoints that were discarded by Confidence-Thresholding, and the corresponding skeleton connections are shown as dashed dark green lines to indicate removed links after post-processing.

**Figure 8.**
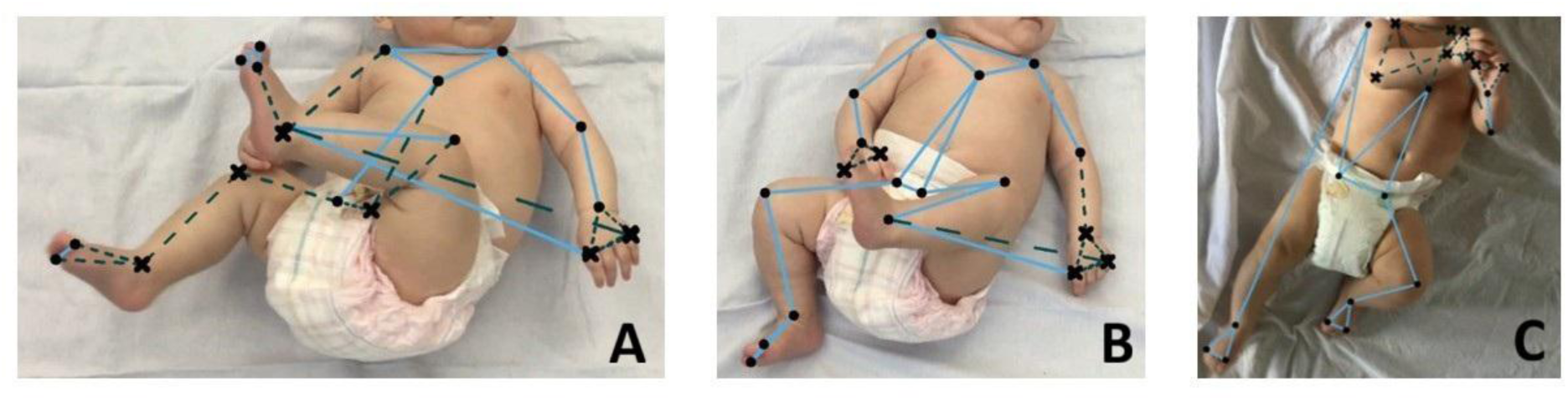
Results after applying the Confidence-Thresholding method with 90% of average confidence threshold.

**Figure 9.**
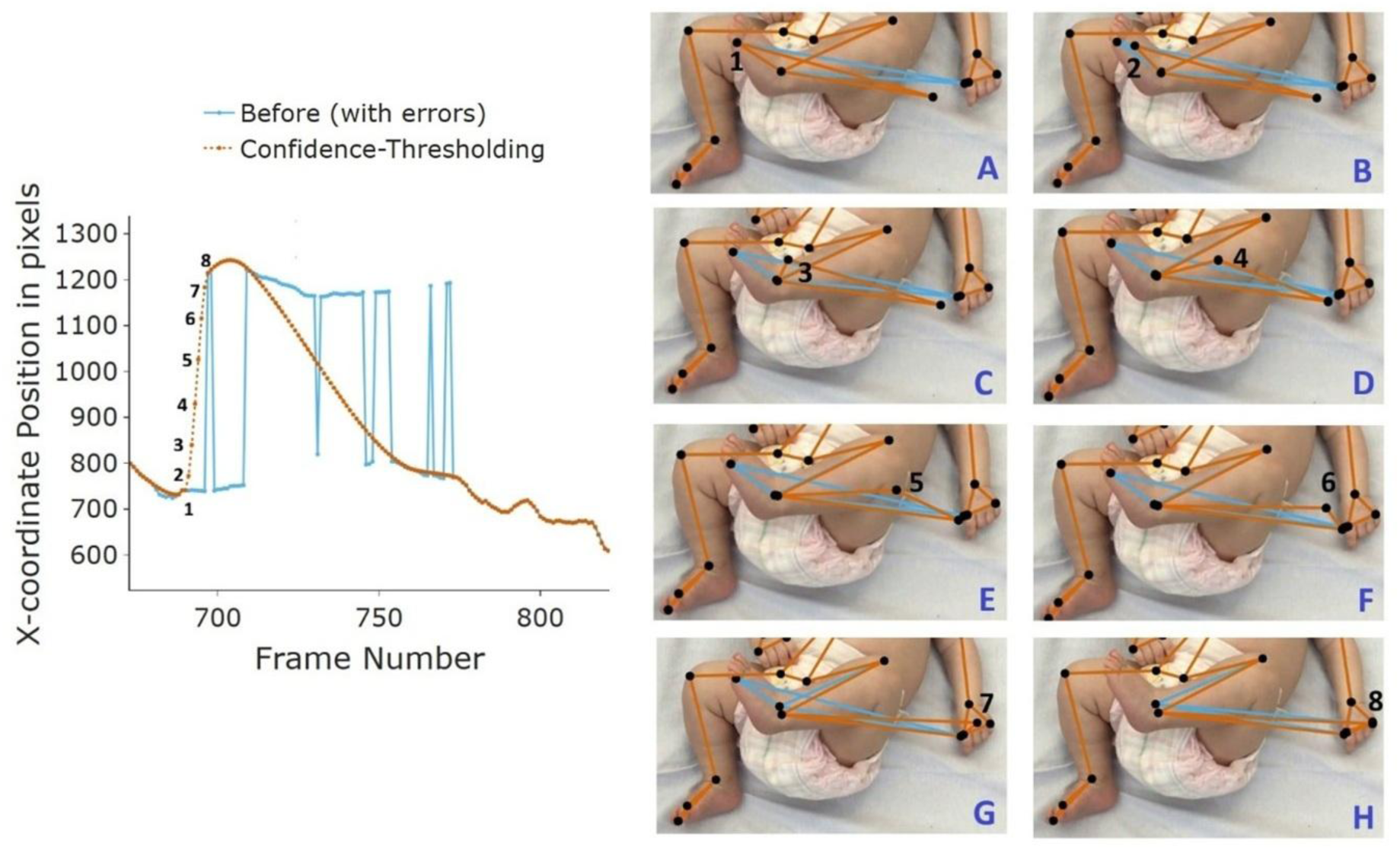
Qualitative results illustrating the Confidence-Thresholding method using the left little toe’ x-coordinate trajectory over time.

**Table 2.**
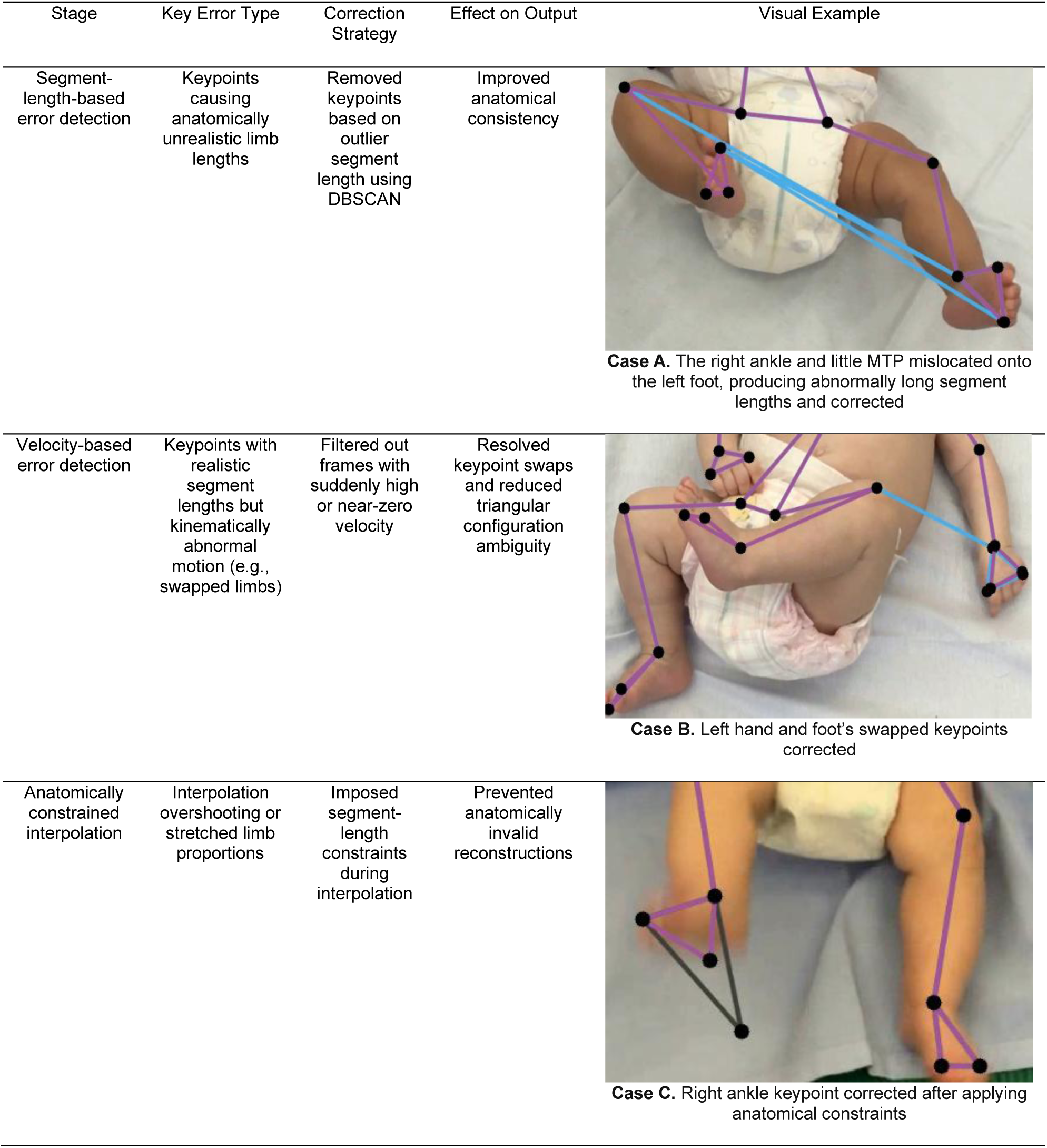
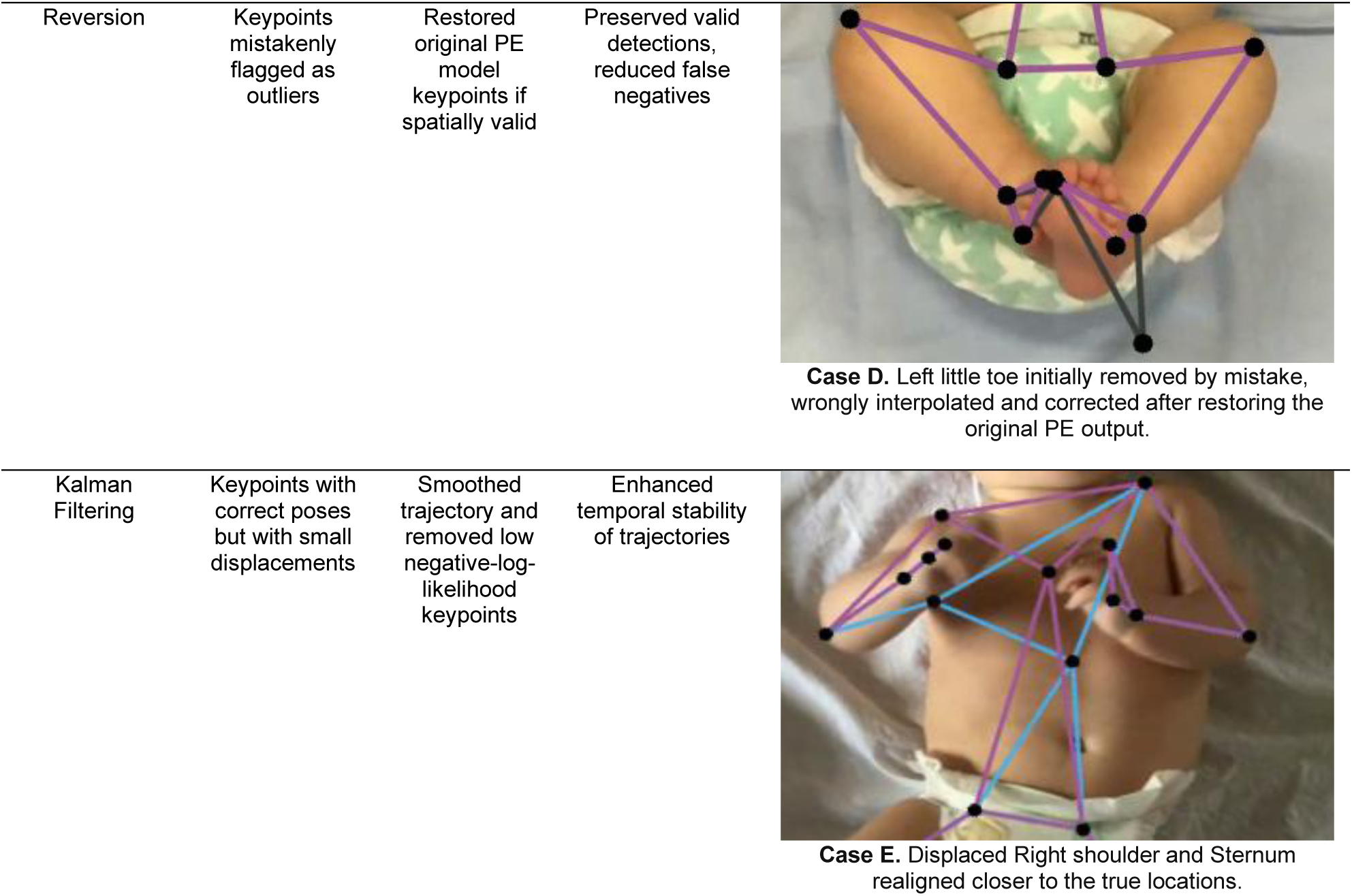
Qualitative assessment results from the CRADLE Method at each stage. *Color codes:* blue lines represent original PE model outputs; orange lines correspond to the Confidence-Thresholding method; gray lines denote intermediate incorrect outputs from the CRADLE pipeline; and purple lines indicate the final output from the CRADLE method.

| Stage | Key Error Type | Correction Strategy | Effect on Output | Visual Example |
| --- | --- | --- | --- | --- |
| Segment-length-based error detection | Keypoints causing anatomically unrealistic limb lengths | Removed keypoints based on outlier segment length using DBSCAN | Improved anatomical consistency | **Case A.** The right ankle and little MTP mislocated onto the left foot, producing abnormally long segment lengths and corrected |
| Velocity-based error detection | Keypoints with realistic segment lengths but kinematically abnormal motion (e.g., swapped limbs) | Filtered out frames with suddenly high or near-zero velocity | Resolved keypoint swaps and reduced triangular configuration ambiguity | **Case B.** Left hand and foot's swapped keypoints corrected |
| Anatomically constrained interpolation | Interpolation overshooting or stretched limb proportions | Imposed segment-length constraints during interpolation | Prevented anatomically invalid reconstructions | **Case C.** Right ankle keypoint corrected after applying anatomical constraints |
| Reversion | Keypoints mistakenly flagged as outliers | Restored original PE model keypoints if spatially valid | Preserved valid detections, reduced false negatives | **Case D.** Left little toe initially removed by mistake, wrongly interpolated and corrected after restoring the original PE output. |
| Kalman Filtering | Keypoints with correct poses but with small displacements | Smoothed trajectory and removed low negative-log-likelihood keypoints | Enhanced temporal stability of trajectories | **Case E.** Displaced Right shoulder and Sternum realigned closer to the true locations. |

In the Confidence-Thresholding method (which ignore keypoints with confidence scores below 90% of their average), several visually accurate keypoints were discarded (indicated by Xs and dark green dashed lines) as shown in Figure 8A (right leg, left hand) and Figure 8B–C (both hands), while some incorrectly localized keypoints with high confidence were retained (e.g., left little MTPs predicted near the left index MCPs in Figure 8A–B, and the right knee in Figure 8C).

Figure 9 provides a detailed example of how interpolation can propagate errors when applied after the Confidence-Thresholding method. The left panel shows the x-coordinate trajectory of the left little toe (numbered point) across several frames. Although the trajectory appears smooth after interpolation, it connects point 1 (a correct keypoint) to point 8 (an unfiltered erroneous keypoint). The right panel (subplots A–H) illustrates this process frame by frame, where the interpolated trajectory gradually shifts the left little toe position toward the incorrect location, resulting in visually incorrect movement.

The CRADLE method employed a multi-stage correction pipeline, with each step targeting a distinct type of error. Table 2 summarizes the correction stages, errors type targeted, the correction strategy used, the effect on output, and representative visual examples. Segment-length-based analysis with DBSCAN identified anatomically incorrect segments and removed corresponding keypoints. For example, in Table 2-Case A, the right ankle and little MTP were mislocated onto the left foot, producing abnormally long segment lengths. Velocity-based anomaly detection with DBSCAN corrected additional errors where segment lengths appeared valid but keypoints were swapped across limbs, such as hand–foot confusions, Table 2-Case B.

Cases C–D in Table 2 shows that subsequent interpolation with anatomical constraints corrected elongated segments, and the reversion step corrected proportion-related distortions and restored valid keypoints that had been mistakenly removed by earlier stages. Finally, Case E (Table 2) shows the realigned right shoulder and sternum to better match anatomical expectations after applying Kalman filter. Collectively, these examples show how successive stages incrementally refine pose trajectories to achieve anatomically consistent outputs.

## 5. Discussion

In this study, we developed a multi-stage post-processing pipeline, CRADLE, to refine 2D video-based infant pose trajectories predicted by a PE model. We evaluated our pipeline’s effectiveness against the conventional Confidence-Thresholding method both quantitatively, using MAE, ΔMAE, aPCK, and net keypoint correction rate, and qualitatively through detailed visual inspections of overlaid pose trajectories. Across all evaluations, the CRADLE method consistently outperformed Confidence-Thresholding approach, demonstrating superior accuracy, anatomical validity, and temporal consistency.

Accurate pose tracking is essential for downstream clinical applications such as GMA, where subtle movement and rotational variations in hands and feet serve as key indicators of neurological risks. However, infants’ distal joints, such as MCPs and MTPs, are highly mobile, prone to occlusion, and therefore frequently mislocalized by automated PE systems. To detect and correct erroneous keypoints, previous studies have implemented the Confidence-Thresholding method. We demonstrated in our results (Figures 8 and 9) that this traditional approach exhibits two key limitations for these distal joints (e.g., MCPs and MTPs) due to its heavy reliance on the confidence scores from the PE model.

First, as shown in Figure 8, the Confidence-Thresholding method can unnecessarily remove accurately predicted keypoints with low confidence, often resulting from occlusion, rapid movement, or complex postures. This may create long temporal gaps in the trajectories, sometimes lasting several seconds, during which meaningful movement information is lost. Furthermore, without sufficient reference points, interpolation over these gaps can produce trajectories that diverge from the true motion path (see Figure 9). Second, mislocalized keypoints that are assigned high confidence, such as MCPs and MTPs during flexed or curled limb postures, remain undetected, as in Figure 8A–C. Moreover, interpolation over partially corrected trajectories can generate smooth but anatomically inconsistent transitions between valid and erroneous keypoints (see Figure 9, left side). Such inaccuracies can, in turn, propagate into automated GMA and other downstream analysis, which critically depend on the precision of pose-derived motion features, potentially compromising their reliability. The CRADLE pipeline, which is independent of confidence scores, addresses these limitations by integrating anatomical features (segment lengths), temporal patterns (velocity-based anomalies), and logical rules based on joint connectivity. The CRADLE approach selectively identifies and corrects erroneous keypoints, while preserving valid predictions, thereby minimising disruption to original trajectories and maintaining the natural continuity of infant movements. Methodological insights for each stage of the pipeline are detailed in the subsequent section of the discussion.

The quantitative evaluation demonstrates the robustness of the CRADLE pipeline. All metrics consistently converged on the same conclusion: the CRADLE method yielded more accurate, temporally stable, and anatomically consistent pose trajectories than the un-post-processed data or the Confidence-Thresholding method. Across all infants and keypoints, MAE was consistently reduced compared with the Confidence-Thresholding method (Figure 3). These results were further supported by the ΔMAE values (Figure 4), which were near zero or positive for nearly all keypoints, indicating that the CRADLE pipeline reduced the distance between corrected and true keypoints, thereby decreasing the MAE after post-processing. Frame-level ΔMAE distributions (Figure 5) further showed a higher frequency of large-magnitude improvements with CRADLE compared with Confidence-Thresholding. Moreover, the relationship between MAE before post-processing and ΔMAE (Figure 6) indicates that the extent of correction was aligned with the severity of the PE prediction error. Rather than uniformly altering trajectories, the CRADLE prioritizes and corrects the erroneous keypoints while preserving those that are already well localized. Together, these findings suggest that the CRADLE pipeline refines trajectories in a controlled and targeted manner, supporting both accuracy and temporal consistency.

Similarly, aPCK values at multiple thresholds (Figure 7) showed that the CRADLE pipeline achieved performance comparable to or better than the original predictions, even under the strictest 5% threshold. Although aPCK averages may not capture localized, frame- and keypoint-level corrections, their stability indicates that the CRADLE method preserves correct keypoint predictions while selectively correcting errors. In contrast, Confidence - Thresholding consistently reduced aPCK across several keypoints, reflecting substantial degradation of localization accuracy across many keypoints. The net keypoint correction rate (Table 1) further highlighted this balance between correcting errors and introducing new ones. Positive values across most keypoints confirmed that the CRADLE method corrected more errors than it introduced, whereas Confidence-Thresholding often yielded negative values due to the removal of visually correct but low-confidence keypoints, which created extended gaps that interpolation then misinterpreted as valid trajectories.

The analysis of challenging cases further highlighted the strengths of the CRADLE method. For example, in Infant #2 (Figure 3), several keypoints of the right hand, including the right little MCP, were only partially annotated in the PE model’s training set due to poor visibility. This lack of annotation limited the model’s ability to learn reliable patterns, resulting in sequences with short alternating sequences of correct and incorrect predictions. In such cases, due to parameter sensitivity, DBSCAN in Stage 1 occasionally misclassifies brief stretches of correct points embedded within longer noisy sequences, leading to their removal. Although this created larger gaps, the anatomically constrained interpolation stage reconstructed trajectories that, while not entirely accurate, remained close to anatomically valid positions and prevented extreme errors. This behaviour was reflected in the aPCK results for Infant #2 (marked * in Figure 7), where Confidence-Thresholding slightly outperformed the CRADLE method for some keypoints on the right hand. However, the MAE results (Figure 3, Infant #2) showed that the CRADLE method ultimately achieved lower error and provided more stable keypoint location than Confidence-Thresholding, even under these complex cases. Overall, these findings highlight the potential of our CRADLE post-processing method for real-world applications where both accuracy and stability are critical.

### 5.1. Methodological Insights

The multi-stage pipeline was designed to address distinct types of keypoint errors while preserving infants’ natural movement patterns, with each stage contributing complementary corrections. Notably, segment lengths emerged as a reliable primary feature for DBSCAN to identify anatomically incorrect keypoints for removal. Compared to raw 2D trajectories (x and y coordinates, separately and combined) and joint angles,^19^ which are strongly behaviour-dependent and therefore vary substantially in the course of normal infant movement, segment lengths primarily depend on anatomy and remain relatively stable over time. As a result, trajectories and joint angles were less reliable for isolating keypoint-level inaccuracies, whereas segment lengths provided a more robust and anatomically interpretable representation, enabling clearer distinction between normal and unrealistic poses. This comparison highlights the importance of selecting the right features that capture the structural relationships of the poses (skeleton) rather than relying solely on raw coordinates. DBSCAN, applied to segment lengths, proved effective in identifying keypoint tracking irregularities. By tuning parameters (*ε* and *minpts*) individually for different body regions according to their mobility, the CRADLE method accommodated variability in movement patterns. For example, static regions such as trunk (e.g. shoulders and ASISs) required smaller values (*ε* = 0.25, *minpts* = 20), whereas dynamic regions such as limbs and face were benefited from higher values (limbs: *ε* = 0.6, *minpts* = 20; face: *ε* = 0.55, *minpts* = 40). This highlights a key advantage of DBSCAN over methods such as k-means, which require a predefined number of clusters, as DBSCAN is capable of handling the complex and highly variable infant movement patterns by detecting clusters of arbitrary size and density. Hierarchical logical rules based on anatomical connectivity of segments, as defined by their lengths, and applied in a proximal-to-distal direction, further enhanced correction accuracy. This proximal-to-distal joint tracking strategy proved particularly valuable in resolving ambiguous configurations, such as triangular arrangements of hand and foot keypoints, where it would otherwise be challenging to infer which joints were mislocalized, for example, determining whether both MCPs or the wrist were incorrect. These explicit decision-making rules not only enhanced robustness but also made the CRADLE method generalizable across diverse infant movements.

While Stage 1 effectively removed many anatomically mislocalized keypoints, some errors remained undetected, particularly when distal hand joints (i.e., wrists, MCPs) were incorrectly localized to anatomically similar foot locations (e.g., ankles and MTPs) as shown in Table 2 (Case B). In such cases, segment lengths appeared anatomically valid despite mislocalization. With the largest errors already removed, at this point, abrupt velocity changes became more apparent, enabling DBSCAN clustering on keypoint velocity to detect these subtle errors. Velocity spikes appeared only at the boundaries of sequences of incorrect keypoint localization, with normal velocities in between. A sliding-window ensured that these temporally consistent but erroneous keypoint positions were also flagged, allowing more complete error detection.

Anatomically constrained interpolation addressed a common yet often overlooked issue in PE post-processing, i.e. trajectory distortions arising from large interpolation gaps as shown in Figure 9. Among the implemented interpolation methods (Modified Akima, polynomial, and cubic splines), Modified Akima interpolation outperformed polynomial and cubic spline based on both visual inspection and aPCK-based quantitative evaluation. By applying segment-length thresholds derived from predominantly correct pre-interpolation data, the CRADLE method maintained reconstructed poses within anatomically valid proportion limits. Furthermore, logical rules (Step 14, of supporting Table S1) identified distal keypoints (MCP, MTP) that drifted too far from their proximal joints (wrists, ankles) during interpolation, resulting in unrealistically long segments (e.g., wrists–MCPs, and ankles–MTPs). These keypoints were then repositioned using anatomical constraints, which reduced the propagation of interpolation-related errors. This constraint-based adjustment effectively prevents unrealistic limb elongation and preserves anatomically consistent motion patterns for smaller joints (MCPs and MTPs), thereby improving the reliability of trajectories for subsequent kinematic analysis.

Although DBSCAN effectively identified errors across complex infant movement patterns, its sensitivity to parameter selection occasionally resulted in the misclassification of originally correct keypoints, particularly for MCPs and MTPs. Such over-flagging unnecessarily removed valid keypoints, resulting in longer temporal gaps that increased interpolation error (Table 2, Case D). To prevent overcorrection, these falsely flagged keypoints were identified by comparing the segment lengths from intermediate corrected (obtained from anatomically constrained interpolation) and raw PE predictions. When the original predictions satisfied anatomical consistency constraints and did not exhibit abnormal segment-length behaviour, the corresponding keypoints were reverted to their original PE predictions, preserving valid data and maintaining the continuity of the true trajectories. This reversion yielded a notable improvement in the overall performance of the CRADLE method, as reflected in Figure 4, where the 95% CI of ΔMAE remained positive or close to zero relative to the Confidence-Thresholding.

In the final stage, we applied the Modified Akima interpolation without any anatomical constraints, due to the nature of the gaps, which were short (≈5–6 frames), after applying the Kalman filter. Because these gaps were bounded by reliable keypoints, unconstrained interpolation effectively restored the correct motion trajectory without risk of anatomical inaccuracy.

## 6. Limitations

We acknowledge that the CRADLE pipeline may have several limitations. First, it may require dataset-specific tuning of DBSCAN parameters, particularly when keypoint motion speeds vary across infant videos. Second, its effectiveness depends on the quality of the initial predictions: when a large proportion of keypoints are incorrectly localized with only sparse intermediate correct predictions, even the reversion step may be insufficient to fully restore trajectory accuracy. As a result, extremely noisy or sparse pose estimates, particularly those produced by models trained on limited or non-diverse datasets, may reduce the reliability of proposed corrections. Third, CRADLE was evaluated using trajectories obtained from a single PE platform (i.e. DLC). Different PE platforms may produce distinct error types, which could influence the effectiveness of the proposed correction strategy. Although test videos encompassed a range of challenging recording conditions (e.g., variations in skin tone, background clutter, and camera motion), further validation across larger datasets and alternative PE platforms is required to assess the generalizability of the CRADLE method.

## 7. Conclusion

We present a multi-stage post-processing pipeline, CRADLE, that effectively refines the quality of 2D infant pose trajectories, particularly in anatomically and clinically critical joints such as the hands and feet. By leveraging spatial and temporal context rather than relying on confidence scores, CRADLE selectively corrects errors while preserving accurate predictions, producing anatomically consistent and temporally stable trajectories. These improvements enhance the reliability of automated infant movement assessments, supporting early detection of neurodevelopmental disorders, and providing a stronger foundation for downstream clinical analyses. Our approach offers a practical, robust solution for improving PE in complex infant movements, with potential impact for both research and clinical applications.

- The study was supported by Friedlander Foundation grant (3720759), through Professor Thor Besier (Auckland Bioengineering Insitute, University of Auckland, New Zealand).
- The authors gratefully acknowledge Nikki Laker and colleagues from Waikato District Health Board, New Zealand for their assistance with participant recruitment and data collection.
- We also acknowledge the use of the eResearch Infrastructure Platform hosted by the Crown comapny, Research and Education Advanced Network New Zealand (REANNZ) Ltd., and funded by the Ministry of Business, Innovation & Employement, URL: https://www.reannz.co.nz, accessed on 17 April 2026).
- We acknowledge that ChatGPT-4o (free version with limited access) was used to refine the manuscript by correcting grammatical errors and enhancing overall readability. All AI-generated suggestions were carefully reviewed and edited by authors to ensure accuracy, integrity, and originality prior to incoporation into the final manuscript.

## Data Availability Statement

Data is not publicly available due to privacy and ethical restrictions.

Received: ((will be filled in by the editorial staff))

Revised: ((will be filled in by the editorial staff))

Published online: ((will be filled in by the editorial staff))

## Supporting Information

Supporting Information is available from the Wiley Online Library or from the author upon request.

