## Supporting material Tables S1, S3, S4, S5, Supporting material Figure S2 for "CRADLE: A Clinically Robust, Anatomy-Aware Post-Processing Framework for Infant GMA Landmark Tracking in 2D Videos"

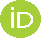

[This Photo](https://www.scirp.org/journal/paperinformation.aspx?paperid=97663) by Unknown Author is licensed under [CC BY-NC](https://creativecommons.org/licenses/by-nc/3.0/)

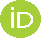

[This Photo](https://www.scirp.org/journal/paperinformation.aspx?paperid=97663) by Unknown Author is licensed under [CC BY-NC](https://creativecommons.org/licenses/by-nc/3.0/)

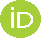

[This Photo](https://www.scirp.org/journal/paperinformation.aspx?paperid=97663) by Unknown Author is licensed under [CC BY-NC](https://creativecommons.org/licenses/by-nc/3.0/)

Manpreet Kaur, Angus J.C. McMorland

School of Exercise, Sports and Rehabilitation Sciences, Faculty of Science, University of Auckland, Auckland,1023, New Zealand;

Hamid Abbasi

Auckland Bioengineering Institute (ABI), University of Auckland, Auckland, 1010, New Zealand;

Centre for Brain Research, University of Auckland, Auckland, 1023, New Zealand;

Department of Physiology, Faculty of Medical and Health Sciences, University of Auckland, Auckland, 1023, New Zealand;

**Table S1.** Pseudocode of the CRADLE pipeline

Algorithm 1: Pose Correction Pipeline

***Input:*** 2D poses predicted by pose estimation (PE) model (24 anatomical keypoints per frame)

***Output:*** Corrected 2D keypoint trajectories

**Stage 1: Segment Length-Based Error Detection**

1. Load PE predicted 2D keypoints.
2. Compute 29 segment lengths per frame between anatomically adjacent keypoints (segments indexed 1–29; see Fig. 1).
3. For each segment time series, apply DBSCAN to classify each segment length as: C (Correct, belonging to a dense cluster) or I (Incorrect, outliers).
4. Store segment-wise correctness labels for subsequent inference.

**Stage 2: Logical Inference and Removal of Incorrect Keypoints**

5. Infer incorrect keypoints using hierarchical logical rules based on connected segment labels.:

Refer to segment numbers 1-29 as labeled in Figure 1.

- **Facial Keypoints**

If (1 & 2 == I): Rt_eye

If (1 & 3 == I): Lt_eye

If (2 & 3 == I): Nose

- **Sternum**

If (10 & 11 == I) or (12 & 13 == I): Sternum

- **Right Upper Limb**

If (4 & 10 == I) or (5 & 4 == I) or (5 & 10 == I): Rt_shoulder

If (4 & 10 == C & 5 == I) or (7 & 9 == C & 6 == I): Rt_elbow

If [(6 & (9 or 7) == I)] or [(5 & 8 == C) & (6 & (9 or 7) == I)]: Rt_wrist

If (7 & 8 == I) or (6 == C & 7 == I): Rt_little_MCP

If (8 & 9 == I) or (6 == C & 9 == I): Rt_index_MCP

- **Special Cases (Right Hand)**
- Two keypoints on the same hand/feet are simultaneously incorrect

If (6 == C) & (9 & 7 == I): Both Rt_MCPs

If (5 == C) & (6 & 7 & 8 == I): Rt_little_MCP and Rt_wrist

If (5 == C) & (6 & 9 & 8 == I): Rt_index_MCP and Rt_wrist

- All three keypoints on the same hand/feet are incorrect

If (6 & 9 & 7 == I): Both Rt_MCPs and Rt_writs

6. Apply symmetric rules to left-upper limb

7. Apply analogous rules for lower limbs using:

Shoulder 🡪 ASIS, Elbow 🡪 Knee, Wrist 🡪 Ankle,

Index MCP🡪 Big Toe, Little MCP🡪 Little Toe

8. Replace all inferred keypoints with **NaN.**

**Stage 3: Velocity-Based Error Detection**

9. For distal keypoints (wrists, ankles, MCPs and MTPs), compute frame-to-frame velocities.
10. Apply DBSCAN to velocity time series to flag abnormal motion.
11. For each flagged keypoint, examine ±2-frame temporal window:

- If velocity ≈ 0, extend incorrect flag to neighbouring keypoints.

12. Replace flagged keypoints with NAN.

**Stage 4: Anatomically-Constrained Interpolation**

13. Interpolate missing keypoints using Modified Akima interpolation.
14. Correct Wrongly interpolated hands and feet:
a. Compute 99^th^ percentile segment length thresholds from Step 5.
b. For segments (7,8,9), if interpolated segment length > threshold:

- If 7 & 8 exceed: correct Rt_little_MCP
- If 8 & 9 exceed: correct Rt_index_MCP
- If 7 & 9 exceed: correct both Rt_MCPs

c. Repeat for left hand and both feet.

**Stage 5: Reversion of Misclassified Keypoints**

15. Revert interpolated keypoints to original PE predictions:
a. If interpolated keypoints are within 10% of max torso length from the original 🡪 keep the original.
b. If segment lengths were correct in original but incorrect after interpolation 🡪 restore original

**Stage 6: Kalman Filter-Based Refinement**

16. Apply forward Kalman filter to each keypoint trajectory and compute negative log-likelihood.
17. Replace keypoints with log-likelihood < (-25) by NaN.
18. Apply the final Modified Akima interpolation to fill gaps.

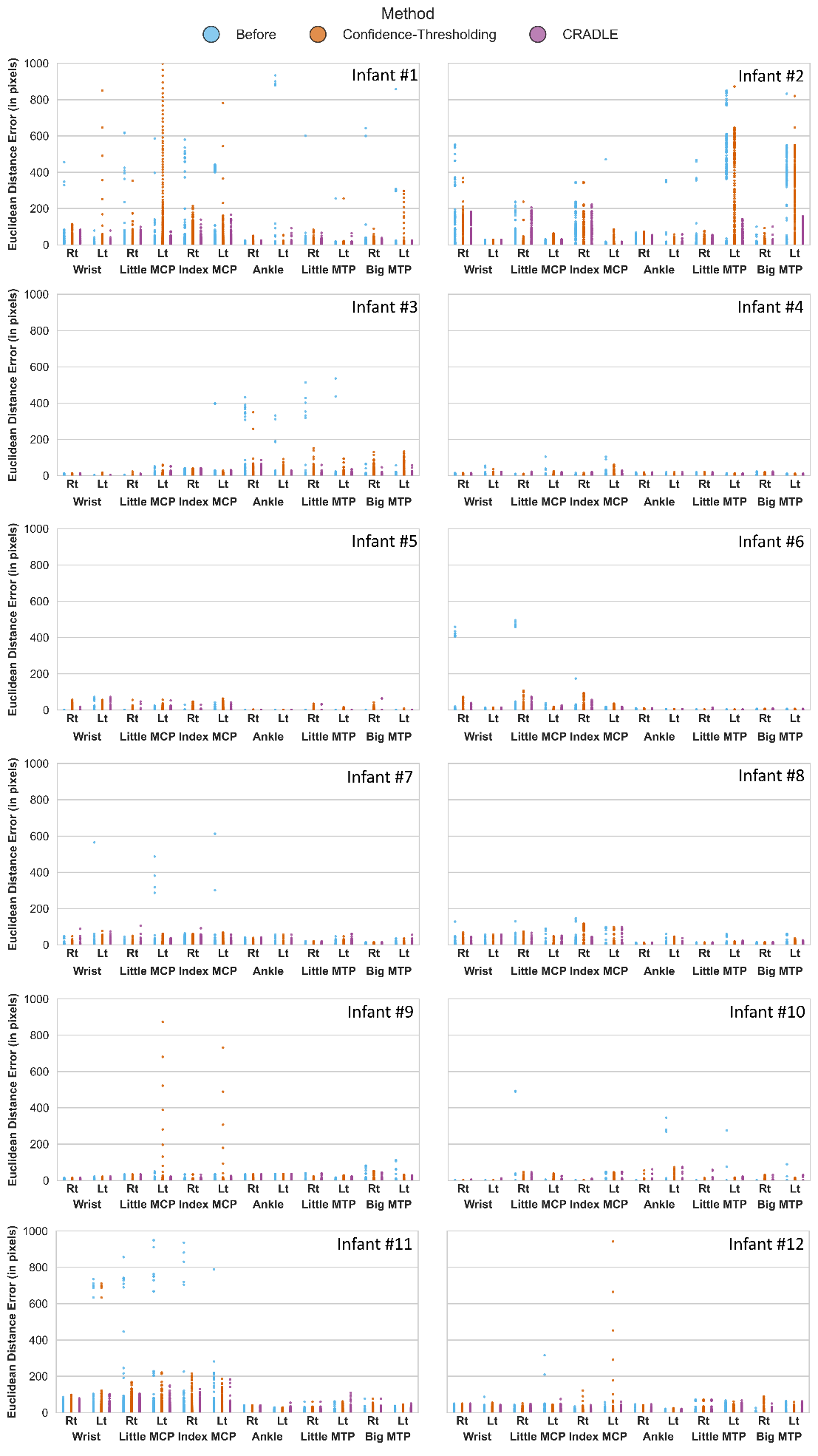

**Figure S2.** Mean absolute error (MAE) distributions for 12 infants. Blue, orange, and purple markers indicate the Before, Confidence-Thresholding, and CRADLE methods, respectively. Each plot shows per-frame MAE (y-axis) for individual keypoints (x-axis) and each dot corresponds to a single frame.

**Table S3.** Averaged across all 12 infants, ΔMAE (pre- minus post- processing MAE) and SD

|  | Confidence-Thresholding | | CRADLE | |
| --- | --- | --- | --- | --- |
| Keypoints | ΔMAE [pixels] | SD | ΔMAE [pixels] | SD |
| Rt_wrist | 0.517 | 1.128 | 1.128 | 5.882 |
| Rt_little_MCP | 0.319 | 0.845 | 0.845 | 9.994 |
| Rt_index_MCP | -1.042 | 0.512 | 0.512 | 1.970 |
| Rt_ankle | -0.263 | 0.122 | 0.122 | 0.452 |
| Rt_little_toe | -0.028 | 0.235 | 0.235 | 3.335 |
| Rt_big_toe | -0.426 | 0.035 | 0.035 | -0.304 |
| Lt_wrist | -8.597 | 0.199 | 0.199 | -90.896 |
| Lt_little_MCP | -4.275 | 0.499 | 0.499 | -27.165 |
| Lt_index_MCP | -4202.175 | 1.155 | 1.155 | -23827.052 |
| Lt_ankle | 0.102 | 0.358 | 0.358 | 6.161 |
| Lt_little_toe | -63.321 | 4.721 | 4.721 | -524.888 |
| Lt_big_toe | -1.426 | 5.013 | 5.013 | -0.646 |

**Table S4.** Average aPCK before, and after post-processing (Confidence-Thresholding and CRADLE)

|  | Before | | | After Confidence-Thresholding | | | After CRADLE | | |
| --- | --- | --- | --- | --- | --- | --- | --- | --- | --- |
| Keypoints | 5% | 7% | 10% | 5% | 7% | 10% | 5% | 7% | 10% |
| Rt_wrist | 0.979 | 0.984 | 0.988 | 0.966 | 0.976 | 0.985 | 0.979 | 0.984 | 0.988 |
| Rt_little_MCP | 0.971 | 0.980 | 0.986 | 0.954 | 0.972 | 0.985 | 0.969 | 0.978 | 0.985 |
| Rt_index_MCP | 0.957 | 0.970 | 0.981 | 0.939 | 0.956 | 0.967 | 0.955 | 0.969 | 0.980 |
| Rt_ankle | 0.983 | 0.990 | 0.995 | 0.974 | 0.984 | 0.992 | 0.983 | 0.990 | 0.995 |
| Rt_little_toe | 0.988 | 0.994 | 0.997 | 0.983 | 0.992 | 0.996 | 0.987 | 0.994 | 0.997 |
| Rt_big_toe | 0.993 | 0.997 | 0.998 | 0.982 | 0.989 | 0.995 | 0.992 | 0.996 | 0.998 |
| Lt_wrist | 0.987 | 0.992 | 0.996 | 0.972 | 0.982 | 0.991 | 0.985 | 0.991 | 0.995 |
| Lt_little_MCP | 0.976 | 0.986 | 0.993 | 0.953 | 0.968 | 0.981 | 0.976 | 0.986 | 0.994 |
| Lt_index_MCP | 0.967 | 0.982 | 0.990 | 0.939 | 0.958 | 0.973 | 0.967 | 0.982 | 0.992 |
| Lt_ankle | 0.989 | 0.992 | 0.995 | 0.983 | 0.988 | 0.993 | 0.988 | 0.993 | 0.996 |
| Lt_little_toe | 0.978 | 0.984 | 0.988 | 0.970 | 0.980 | 0.985 | 0.981 | 0.989 | 0.993 |
| Lt_big_toe | 0.970 | 0.977 | 0.982 | 0.956 | 0.967 | 0.972 | 0.970 | 0.979 | 0.986 |

**Table S5.** Net Correction Rate per 10k frames

|  | Confidence-Thresholding | | | CRADLE | | |
| --- | --- | --- | --- | --- | --- | --- |
| Keypoint | 5% | 7% | 10% | 5% | 7% | 10% |
| Rt wrist | -188 | -122 | -43 | -7 | -7 | 2 |
| Rt hand little finger MCP | -150 | -75 | -22 | -16 | -13 | -9 |
| Rt hand index finger MCP | -191 | -133 | -120 | -12 | -9 | 2 |
| Rt ankle | -86 | -55 | -29 | -1 | 1 | 2 |
| Rt little toe | -57 | -25 | -9 | -7 | -1 | 3 |
| Rt big toe | -127 | -79 | -36 | -8 | -1 | 2 |
| Lt wrist | -185 | -137 | -64 | -12 | -8 | -7 |
| Lt hand little finger MCP | -205 | -159 | -107 | -3 | 1 | 9 |
| Lt hand index finger MCP | -259 | -202 | -145 | 3 | 13 | 21 |
| Lt ankle | -56 | -39 | -19 | -1 | 3 | 5 |
| Lt little toe | -52 | -27 | -11 | 13 | 23 | 30 |
| Lt big toe | -105 | -80 | -67 | 2 | 9 | 20 |
